# Assessing and Optimizing Low-Frequency Somatic Mutation Detection: A Multi-Platform High-Throughput Sequencing Perspective

**DOI:** 10.64898/2026.05.28.728367

**Authors:** Baosheng Feng, Yixin Lin, Lei Liu, Qunting Lin, Yuan Lin, Yongfang Liu, Jiatao Li, Chaoyu Lei, Chang Chen, Manman Yang, Xuejie Peng, Zhiliang Zhou, Qin Yan, Lei Sun, Qigang Li

## Abstract

The availability of multiple commercial short-read sequencing platforms necessitates systematic cross-platform performance comparisons, particularly for challenging applications such as low-frequency somatic mutation detection. Here, a large-scale targeted sequencing dataset from five Genome in a Bottle (GIAB) human genomic DNA reference standards, HG001 to HG005, alongside Twist Biosciences cfDNA reference standards featuring 1% variant allele frequency (VAF), was generated by six platforms (NovaSeq 6000, NovaSeq X, FASTASeq 300, GenoLab M, SURFSeq 5000, and MGISEQ-T7). To build a realistic benchmark while keeping authentic sequencing backgrounds, we developed PosMix, a simulating tool that generates position-specific VAFs. To overcome the limitations of conventional variant callers (high recall with poor precision for VarScan2, higher precision with lower recall for Strelka2/Mutect2), we developed SomaticXGB, a machine learning-based caller. In this study, SURFSeq 5000 consistently exhibited the lowest error rates and achieved superior accuracy for VAFs as low as 0.5%, outperforming all other sequencing platforms. On the other hand, SomaticXGB attained F1 scores of approximately 0.92 on simulated datasets with VAFs ranging from 0.5% to 1.5% and 0.89 on Twist 1% standards, substantially outperforming conventional methods. This work delivers a valuable rich multi-platform data resource, offering a standardized pipeline for performance benchmarking and a machine learning-based strategy for optimized somatic mutation detection.

## Introduction

The emergence of high-throughput sequencing platforms, particularly short-read sequencers characterized by ultra-high data output and low cost, has significantly improved our ability to detect somatic mutations, which may played a key role in cancer research, diagnosis, and therapy[1–3]. Over the past decade, Illumina and its competitors, such as MGI, GeneMind, and Element Biosciences, have introduced a range of short-read sequencers, making the toolkit accessible to researchers in biology and medicine. Therefore, it is necessary to develop robust approaches that work across platforms for somatic mutation detection and performance evaluation.

However, building a system for performance assessment and method optimization is challenging, particularly for somatic mutations with low variant allele frequencies (VAFs). Detecting somatic variants is affected by several factors, including tumor heterogeneity, sample type (formalin-fixed, paraffin-embedded or fresh frozen), library preparation, sequencing platform and data interpretation, especially when dealing with low-frequency variants, particularly those with a VAF below 2%[4,5]. Specifically, during library preparation, cytosine deamination and guanine oxidation can elevate the occurrence of low-frequency CT/GA and GT/CA variants, respectively[6,7]. Although errors from library preparation and sequencing could be corrected by Unique Molecular Identifiers (UMIs) technique, allowing reliable detection of mutations as low as 0.5%[8,9], the UMI-based correction process require an ultra-high sequencing depth (e.g. 10,000X to 30,000X) and an additional data analysis workflow, making it less cost-effective. Moreover, using UMIs for error correction reduces performance discrepancies among platforms, which would confound our cross-platform comparison. In this study, a single library was sequenced on six compatible sequencing platforms and data analysis was performed without UMI, allowing us to focus on evaluating how different sequencing platforms affect the detection of low-frequency somatic mutations.

Additionally, commercial standard tumor cell lines or reference DNA materials often contain a defined list of true somatic variants, but whether a detected variant not listed is a false positive remains ambiguous. For instance, in the widely used HCC1395/HCC1395BL tumor/normal cell line pair, only 20% of variants in the given list are low-frequency (VAF < 5%)[10]. But actually, tumor cell lines harbor a large number of low-frequency somatic mutations[11,12]. Therefore, we infer that the mutation set of HCC1395/HCC1395BL tumor/normal cell line pair is incomplete, making it difficult to evaluate recall and precision metrics of sequencing platforms with high confidence. In addition to constructing reference DNA samples from tumor/normal cell line pair, *in-silico* simulation is another strategy to generate data sets for performance benchmarking[13]. A commonly used method involves mixing data from one genome (donor, simulated tumor origin) into another genome (background) at a certain ratio[14]. However, this approach could incorporate all variants of the donor genome and assign them with certain frequencies in the simulated genome, which does not align with the actual situation where a tumor carries variants with varying frequencies. Another common methods that directly modifies bases, such as BAMSurgeon[15] and VarBen[16], do not fully represent real sequencing outcomes, for example, by changing an A to a T while keeping the original A’s quality score. To preserve original sequencing results and simulate somatic mutations at specified frequencies at specified genomic positions, we developed a tool called PosMix with HTSlib[17], and used it to generate synthetic tumor-normal pair data sets with a complete set of somatic mutations.

In this study, we sequenced 5 reference samples HG001-HG005 on six different platforms (Supplementary table 1). Using PosMix, we generated synthetic tumor data set to evaluate performance and to develop data analysis method (Supplementary table 2). We assessed the platforms with established tools including Mutect2[18,19], Strelka2[20] and VarScan2[21], and we also developed a novel caller named SomaticXGB for somatic mutation detection. Our results showed that instruments with lower sequencing error rates, such as SURFSeq 5000 from GeneMind, achieved higher accuracy in detecting low-frequency mutations. SomaticXGB outperformed commonly used callers across multiple platforms, which is also confirmed by using synthetic tumor standards from Twist. In summary, our comprehensive experimentation, simulation, validation, benchmarking and optimization offered reliable references for detecting somatic mutations in the “multi-platform era”.

## Results

### PosMix enables simulations of somatic variants with position-specific frequencies

For instance, knowing that at a specific position HG001 is homozygous AA (matching the reference sequence) and HG002 is heterozygous AT, PosMix randomly discards 2% of HG001 (background/normal genome) reads covering this position and replaces them with an equal number of covering reads from HG002 (donor genome), generating a mixed alignment file (BAM format) which consists of all remaining reads of HG001 and the replacing reads from HG002. The resulting BAM file contains the desired mutation at certain position with a expected VAF of 1%, and serves as the tumor component in the normal-tumor pair. Additionally, a subset BAM file of HG001 can be generated to serve as the normal component. To demonstrate the performance of PosMix, we examined 103 simulated variants with specified VAFs ranging from 0.5% to 5%. The calculated VAFs were found to be close with the specified frequencies across various platforms and frequency ranges (Fig. 1b), indicating that PosMix successfully achieved position-specific frequency simulation.

**Fig. 1.**
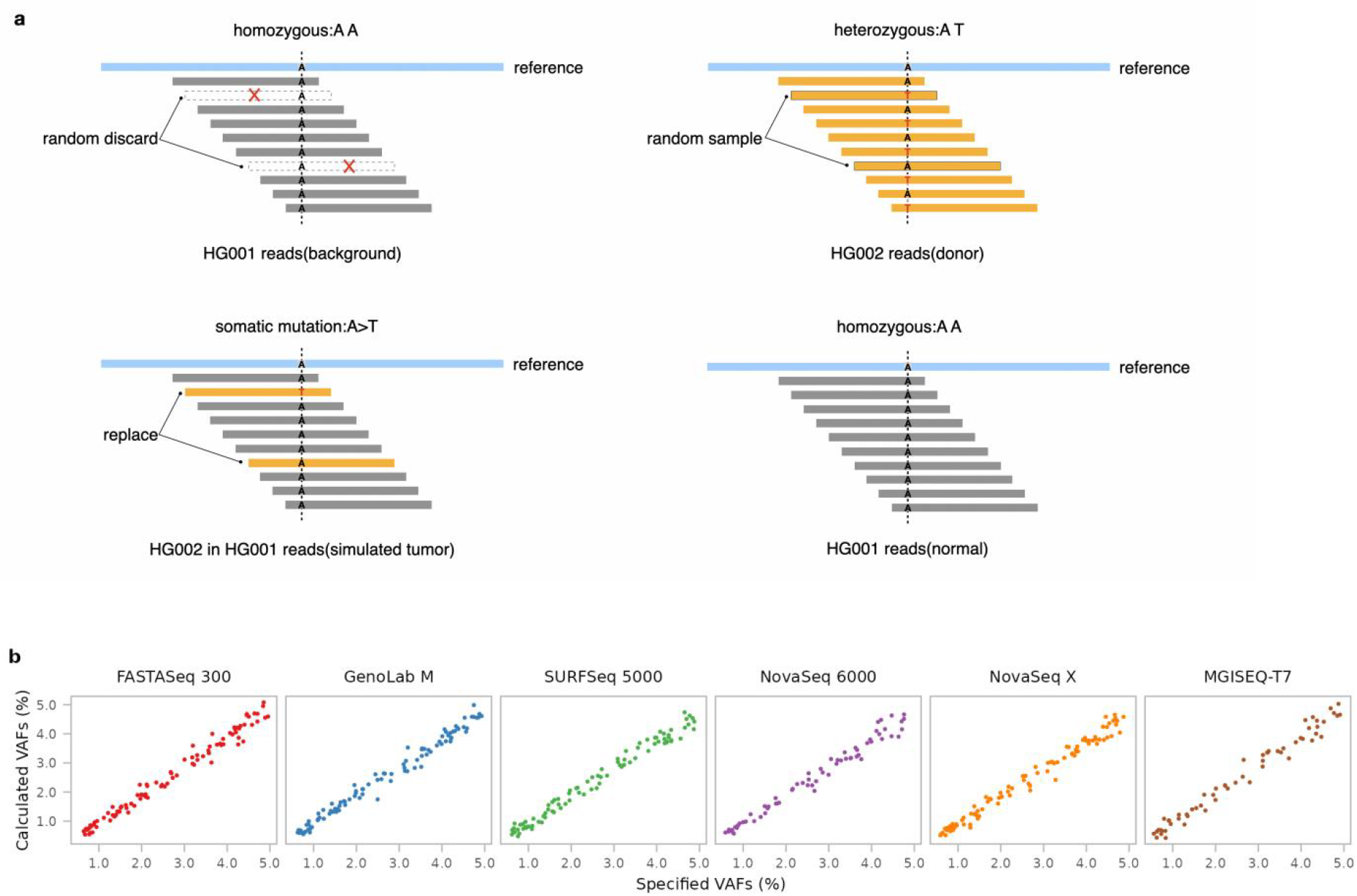
Schematic and validation of PosMix. **a**, Schematic workflow of PosMix, the simulation tool. **b**, Consistency between the specified (expected) and calculated (observed) variant allele frequencies (VAFs) across six sequencing platforms.

### Performance evaluation across multiple platforms using mainstream somatic variant methods

We utilized five reference standards, HG001 to HG005, to construct two types of libraries for sequencing. One type for five sequencing platforms: NovaSeq 6000, and NovaSeq X, FASTASeq 300, GenoLab M, SURFSeq 5000, and the other type for MGISEQ-T7 platform, giving 27 distinct libraries, and 75 sequencing datasets, generating a data collection of 3.22 Terabases (Tb) (Supplementary Table1). Both types of libraries were constructed using iGentech’s OncoPanel T364V1, targeting 2.1 megabase genomic regions encompassing 641 genes.

We examined the performance of six sequencing platforms by mixing HG002 reads into HG001 with PosMix (Supplementary Table 2). In the simulation, there were 414 variants, six low-frequency levels (0.5%, 1%, 1.5%, 2%, 3%, 5%), five average sequencing depths (900X, 1800X, 2700X, 3600X, 4500X). We developed two analyzing methods using three commonly used variant calling tools (callers): “VarScan2_fpfilter” represents the results of the original VarScan2 output after being filtered by its built-in tool, fpfilter; “Strelka2_Mutect2” takes the union of the variants called by Strelka2 and Mutect2. See Materials and Methods for details of the simulation process and development of the two methods.

As expected, higher sequencing depth and higher allele frequency led to better recall, precision and F1 scores across all library types and sequencing platforms. The most significant difference was between the two variant callers. VarScan2_fpfilter maintained a consistently high recall but suffered from significantly low precision while Strelka2_Mutect2 exhibited lower recall but higher precision. This indicates that VarScan2_fpfilter missed fewer true mutations, whereas Strelka2_Mutect2 yielded fewer false positives. For instance, at an VAF of 1% and a sequencing depth of approximately 1800X, the recall for Strelka2_Mutect2 varied between 0.45 and 0.89, whereas VarScan2_fpfilter exhibited a recall between 0.92 and 0.96. In terms of precision, Strelka2_Mutect2 maintained a range of 0.91 to 0.93, whereas VarScan2_fpfilter had a precision range of approximately 0.06 to 0.17 (Fig. 2d, Supplementary Table 3).

**Fig. 2.**
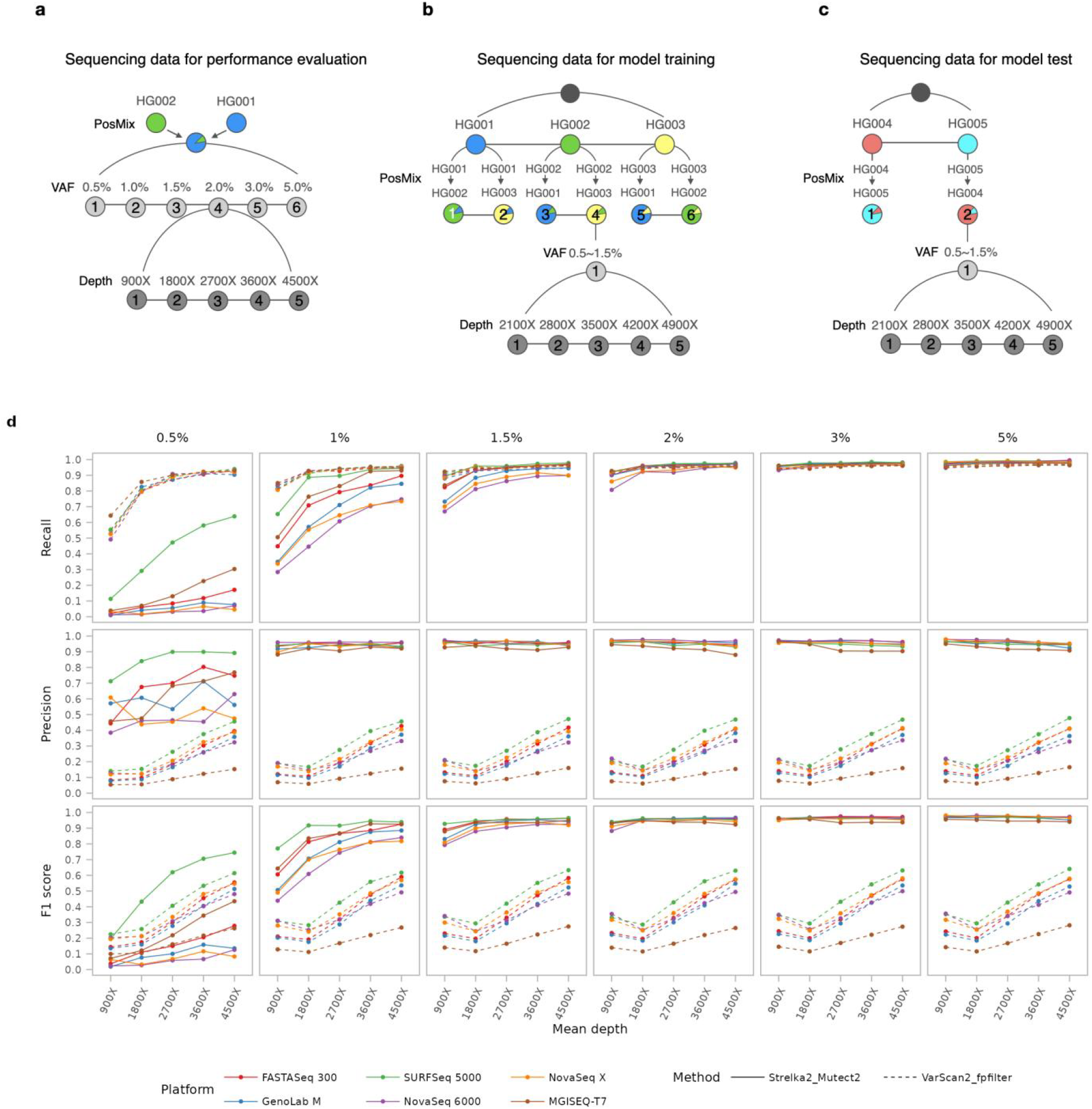
Simulation workflows and platform performance benchmarking. **a-c**, Schematic representation of data utilization for performance evaluation (a), model training (b), and model testing (c). **d**, Comparison of somatic calling metrics (Recall, Precision, and F1 score) across various sequencing platforms, VAF tiers, and sequencing depths. All reported depths represent the raw average coverage before duplicate removal.

Discrepancies among the six platforms were also noted, although these differences were diminished as VAF and/or sequencing depth increased. At the 0.5% VAF by Strelka2_Mutect2, with the increasing of sequencing depth, a marked improvement of recall was observed on the SURFSeq 5000 platform, however, even under high sequencing depth, this detection remained challenging on other platforms, including NovaSeq 6000 and NovaSeq X. Moreover, the SURFSeq 5000 also showed exceptional performance in both precision and F1 score (Fig. 2d). These results suggested that SURFSeq 5000 outperformed other sequencing platforms in detecting somatic variants.

### Superior cross-platform calling performance of our machine learning method

Beyond evaluating established calling methods, we also endeavored to construct a more accurate cross-platform somatic variant caller leveraging machine learning techniques. Given that the raw output from VarScan2 exhibits high recall but also a significant rate of false positives, we aimed to refine these results using machine learning techniques. Initially, with the simulated data described below, we obtained raw variant calls from VarScan2 without applying the fpfilter. Subsequently, we utilized Mutect2’s force-calling method and bam-readcount[22] to extract features for each candidate variant, totaling 17 distinct features (Supplementary Table 4). Then we trained our cross-platform model using XGBoost[23]. Details of the feature engineering and model training methodologies were provided in Materials and Methods.

We employed PosMix to mutually mix sequencing data from HG001, HG002, and HG003 across six platforms. For each pair of sequencing data, we set five distinct sequencing depth levels (2100X, 2800X, 3500X, 4200X, 4900X), yielding a total of 180 simulated normal/tumor pairs of BAM files for model training. Collectively, 2,134 germline variants were used to simulate 64,020 somatic mutations with frequencies randomly assigned between 0.5% and 1.5% across six platforms (Supplementary Table 2).

We employed the same methodology, using PosMix to mix data from HG004 and HG005 to generate simulated data to test our model. This resulted in a total of 25,890 simulated somatic mutations with varying frequencies of 0.5%-1.5%, simulated from 863 distinct germline variants (Supplementary Table 2).

As shown in Fig. 3 and Supplementary Table 5, the results regarding platforms, depths, and methods are consistent with those previously obtained using simulations with HG001 and HG002. SURFSeq 5000 still demonstrates the best overall performance. Likewise, there is a clear disparity between platforms that diminishes as sequencing depth increases, with SURFSeq 5000 outperforming others, particularly at lower depths. Our new caller, named SomaticXGB, maintains the recall of the original results from VarScan2, significantly reduces false positives, and enhances precision and F1 scores, performing strikingly better than the other two methods. For instance, on the NovaSeq X platform at a depth of 2800X, the recall and precision of SomaticXGB (threshold 0.1) were comparable to the best recall and precision of the two other methods, resulting in a F1 score of 0.92, largely outperforming Strelka2_Mutect2’s 0.76 and VarScan_fpfilter’s 0.44. Notably, SomaticXGB markedly improved both performance and stability across all platforms. For example, at the minimum depth of 2100X, the F1 scores for SomaticXGB (threshold 0.1) on all platforms ranged between 0.87 and 0.92, whereas Strelka2_Mutect2 achieved scores from 0.69 to 0.89, and VarScan_fpfilter’s scores spanned from 0.15 to 0.41. This indicates that our model is highly effective and has robust cross-platform applicability.

**Fig. 3.**
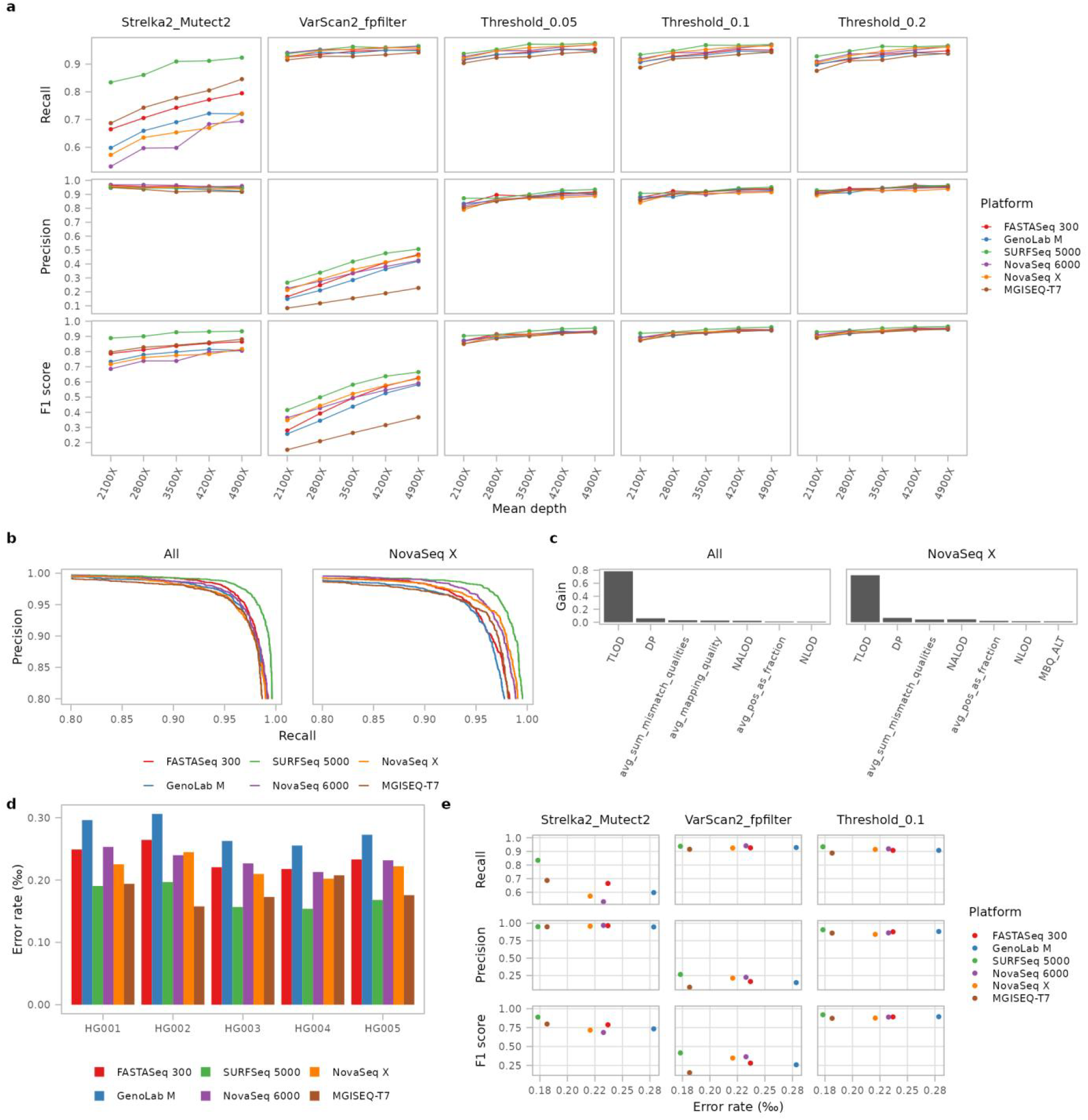
SomaticXGB and SURFSeq 5000 show better performance. **a**, Recall, precision and F1 score across methods, platforms and depths. **b**, Platform receiver operating characteristic (ROC) curves of two models training from all data and only NovaSeq X data. **c**, Comparison of top feature importance between two models. **d**, Error rates of HG001-HG005 across platforms. **e**, Scatter plots of averaged error rates of HG001-HG005 and performance metrics across platforms.

To assess the cross-platform robustness of our approach, we trained a model using only NovaSeq X data and evaluated it on the six platforms. The recall and precision curves for the model trained on the data of all platforms and the model trained solely on NovaSeq X data were nearly close, with both achieving AUC scores above 0.994 (Fig. 3b). At a threshold of 0.1, both models exhibited high performance on multiple platforms in terms of recall, precision, and F1 score (Supplementary Table 6). Furthermore, we noticed a high degree of congruence in feature utilization and weighting between the two models (Fig. 3c and Supplementary Table 7 and 8), indicating that their working mechanisms are strikingly similar. In summary, our machine learning method has successfully identified features that are universally effective across platforms, and these features can substantially improve detection capabilities.

Interestingly, SURFSeq 5000 showed a notably superior performance even on a model trained by data exclusively from NovaSeq X (Fig. 3b and c). Along with other observed performance divergences among platforms, we speculated that this may be related to the sequencing accuracy of each platform. Among five samples across six platforms, SURFSeq 5000 delivered the lowest error rate in four out of five, except for HG002 (Fig. 3d). At sequencing depth of 2100X, SURFSeq 5000, with the lowest error rate, also showed the highest Recall, Precision, and F1 score across all the three methods (Fig. 3e).

### Cross-platform performance validated by Twist Biosciences cfDNA reference standards (VAF of 1%)

In addition to our simulated test sets, we constructed three replicate libraries for both 0% and 1% Twist standard materials to evaluate the performance of four platforms — GenoLab M, SURFSeq 5000, NovaSeq 6000, and NovaSeq X—as well as our new method (Table S9). 1% Twist reference standard includes background DNA derived from human cell-free DNA (cfDNA) and 458 synthetic mimic somatic variants with expected VAFs of 1%, 221 of which are covered by our panel.

Consistent with the aforementioned findings, SomaticXGB outperformed the other two methods overall. For instance, the recall of “Threshold_0.1” was comparable to VarScan2_fpfilter, with both almost exceeding 0.95, while its precision was around 0.5, largely lower than the previously high level above 0.9. Additionally, among the substitution types of false positives, a distinct enrichment was observed for CT/GA and AG/TC transitions (Fig. 4b), which is also observed in the wild type (Fig. 4c). According to results from Twist Biosciences, the v1 used in this study has a higher background mutation rate compared to their optimized v2[24]. We speculated that the decline in performance might be related to the higher non-sequencing noise in the Twist background DNA data, such as cytosine deamination leading to an increase in CT/GA mutations. To reduce background noise, we integrated the 1% Twist variants into HG001 data using PosMix, yielding synthetic tumor data that contains 221 mutations on HG001 background. We found that the precision scores of all three methods were improved; SomaticXGB reached around 0.85. The average F1 score for SomaticXGB across the four platforms was 0.89, significantly higher than the other two methods’ F1 scores — 0.72 for Strelka2_Mutect2 (Wilcoxon test p value = 6.104e-05) and 0.38 for VarScan2_fpfilter (Wilcoxon test p value = 6.104e-05) (Fig. 4d). Therefore, we concluded that background mutations serve as a primary confounding factor in low-frequency mutation detection. The PosMix-based simulation strategy effectively reduced this confounding effect by introducing predefined somatic variants into a well-characterized HG001 background while retaining authentic sequencing errors and experimental noise, enabling a cleaner assessment of platform- and algorithm-dependent detection performance. Concurrently, our machine learning method demonstrated a performance that remains significantly superior to the other two conventional methods.

**Fig. 4.**
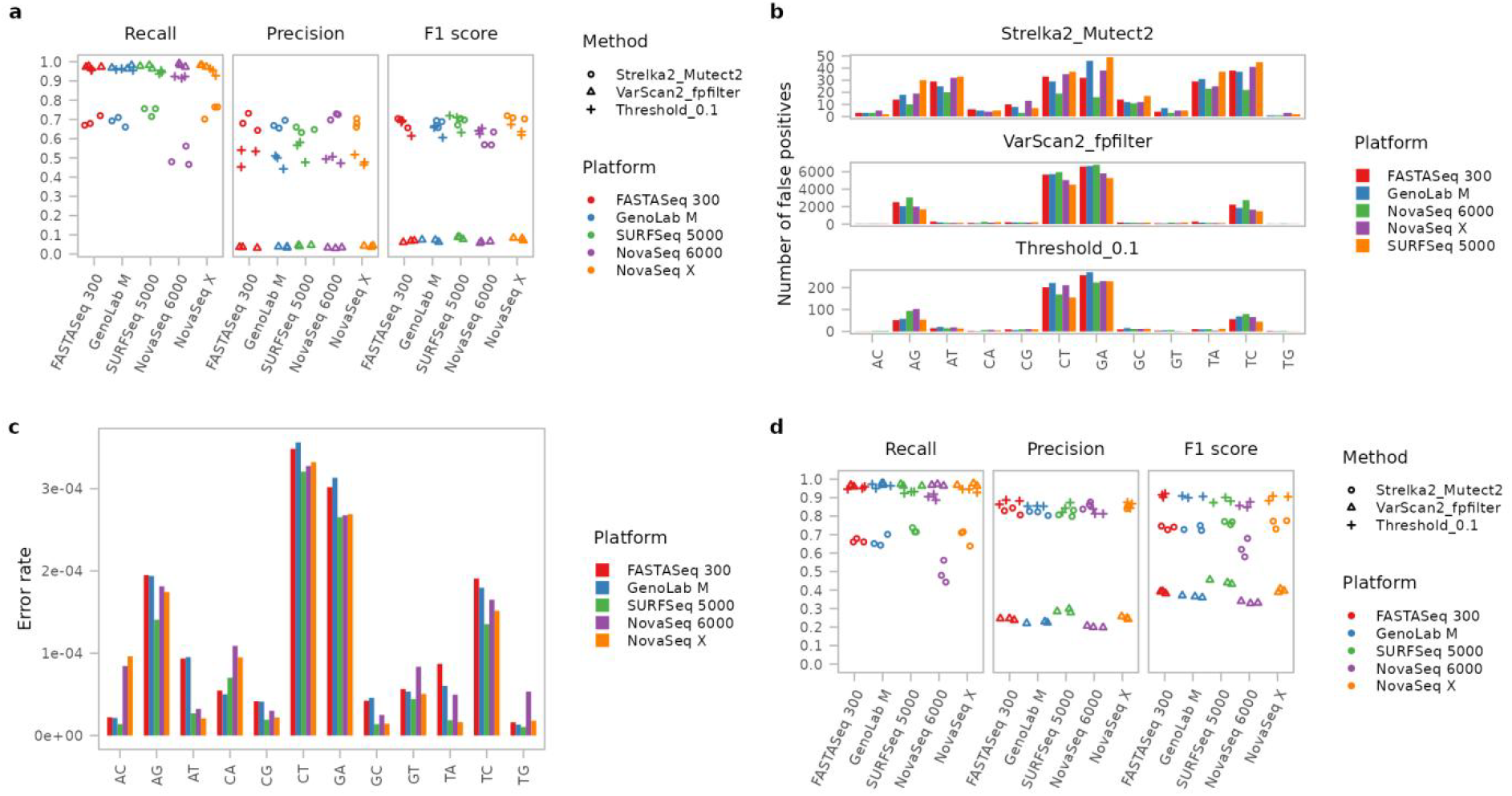
Performance of SomaticXGB validated by Twist DNA standards. **a**, Recall, Precision and F1 score of SomaticXGB and other 2 methods across platforms using 0% and 1% Twist DNA samples. **b**, CT/GA, AG/TC enrichment in the three methods. **c**, High error rates of CT/GA and AG/TC in 0% Twist DNA. **d**, Great increase in precision and F1 metrics when using HG001 as background.

## Discussion

Here, we focused on examining and advancing the performance of various sequencing platforms in detecting low-frequency somatic mutations. We first developed a novel simulation tool that performs position-specific simulation and maintains the authenticity of sequencing data. Many factors in the experimental processes such as DNA standard manufacturing and library preparation can elevate the background mutation rate, such as DNA damage during experiment and replication during PCR reaction. Our simulation method, however, allows us to extract and focus on the differences between platforms. In the future, we could integrate somatic variants derived from tumor cell lines or other tumor reference standards to build more realistic simulation settings.

As previously shown[13,14], it is difficult to achieve high sensitivity (recall) and precision for variants below 2% using conventional callers (e.g., Strelka2, Mutect2, VarScan2). However, our optimized caller, SomaticXGB, can achieve this. We also found substantial differences between platforms, with the SURFSeq 5000—being the most accurate—showing the best overall performance. Remarkably, it achieved at lower sequencing depths the same level of performance that other platforms required higher depths to reach. In summary, the final detection performance of low-frequency variants is by various factors including experimental procedures, sequencing, and analytical methods, and our study provides comprehensive and reliable reference information for the assessment and utilization of short-read sequencing platforms.

## Acknowledgments

This work was supported by the National Science and Technology Major Project of China (Grant No. 2023ZD0505901).

## Materials and Methods

### DNA resources

Germline variants were from HG001, HG002, HG003, HG004 and HG005 bought from Coriell Institute (https://www.coriell.org); the catalog items IDs are NA12878, NA24385, NA24149, NA24143, NA24631, respectively. Synthetic somatic variants from Twist cfDNA Pan-Cancer Reference Standard v1 (https://www.twistbioscience.com) of VAF of 1% and 0% were used to confirm the performance of our new method.

### DNA fragmentation and library preparation

DNA of HG001-HG005 were fragmented. for each library, 250ng were input into ultrasound disruption for 250 seconds in nuclease-free water (UltraPure™ DNase/RNase-Free Distilled Water). IGT® Fast Library Prep Kit v2.0 (C10022) are used for library preparation. Two types of sequencing libraries were constructed: TruSeq and DNBSEQ libraries. The following kits from iGeneTech were used for TruSeq libraries: TargetSep One® Hyb&Wash Kit V2.0 (Module B,66102201), TargetSep Cap Beads&Nuclease-Free Water (C10422), TargetSep One® Hyb&Wash Kit V2.0 (Module A, C10282), TargetSep One® Hyb&Wash Kit (Module C, for Illumina, C10572), TargetSep One® Eco Universal Blocking Oligo (for Illumina, C80502); For DNBSEQ libraries, the following kits from iGeneTech were used: TargetSep One® Hyb&Wash Kit V2.0 (Module B, 66102201), TargetSep Cap Beads&Nuclease-Free Water (C10421), TargetSep One® Hyb&Wash Kit V2.0 (Module A,C10281), TargetSep One® Hyb&Wash Kit (Module C, for MGI DI), TargetSep One® Eco Universal Blocking Oligo (for MGI DI,C80531). TargetSeq® Pan-Cancer Panel (T364V1, PT1004172) from iGeneTech were used to target cancer genes.

### DNA sequencing

TruSeq libraries were sequenced on a FASTASeq 300 instrument (GeneMind) with sequencing kit (v2.0, FCH, 300 cycles), GenoLab M instrument (GeneMind) with sequencing kit (v4.0, FCH, 300 cycles), SURFSeq 5000 instrument (GeneMind) with sequencing kit (v1.0, FCH, 300 cycles), NovaSeq 6000 instrument (Illumina) at 2×150 bases read length and a NovaSeq X instrument (Illumina) at 2 × 150 bases read length. DNBSEQ libraries were sequenced on a MGISEQ-T7 instrument (MGI) at 2×150 bases read length.

### Read processing, alignment and quality assessment

Fastp[25] (v.0.23.4) was conducted with default settings to assess sequencing quality and trim adaptor sequences. Trimmed reads were mapped to the human reference genome GRCh37 using BWA-MEM (v.0.7.17-r1188) with “-K 100000000 -k 32 -M”. Bamdst (v.1.0.7) and Samtools were used to perform post-alignment quality control, including: fraction of PCR duplicate reads, amount of target reads (in panel region), amount of target reads without duplicates, fraction of target reads in all reads, fraction of target data in all data, average depth and average depth without duplicates.

### Development of PosMix

PosMix is a high-performance C++ program that employs HTSlib for reading and manipulating BAM files, which is available in in a GitHub repository (https://github.com/liqg/posmix). The schematic of PosMix is displayed in Fig. 1a.

### Simulation for PosMix validation, evaluation using existing methods, model training and model testing

The gold standard reference variant calls of HG001-HG005 were obtained from Genome in a Bottle (GIAB). Variants located in high-confidence regions, and with a distance of no less than 50 base pairs between any two sites, were utilized for simulation. For PosMix validation, 103 germline variants of HG002 were inserted into HG001 data to create simulated tumor datasets with VAFs ranging from 0.5% to 5%. HG001 and HG002 were used for performance benchmarking using existing somatic variant calling tools. HG001, HG002 and HG003 were used to create simulated data to train our machine learning model; HG004 and HG005 were used for testing the performance of our new method. Using different samples for training and testing allows for an accurate assessment of the model’s performance. Fig. 2a illustrated the simulation schema, and supplementary table 1 described the details of simulation for model training and model testing.

### Somatic SNV calling

We developed two pipelines to call somatic variants using three somatic variant callers, Mutect2 (GATK v4.2.6.1), Strelka2 (v2.9.10) and VarScan2 (v2.4.3). One pipeline called Mutect2_Strelka2 combines results of the two callers. For Mutect2, somatic mutation candidates were called by the default configuration, and then filtered by GATK FilterMutectCalls under the configuration of “--min-reads-per-strand 1 --min-median-base-quality 20 --normal-p-value-threshold 0.01”. Strelka2 was run with the default configuration. The other pipeline called VarScan2_fpfilter. Firstly, samtools mpileup was performed with the configuration of “-d 1000000 -q 30 -Q 30 -A -B -x --excl-flags UNMAP,SECONDARY,QCFAIL,DUP”. Then VarScan2 was performed to analyze paired tumor-normal pileup files under the configuration of “--min-coverage-normal 10 --min-coverage-tumor 100 --min-freq-for-hom 0.75 --normal-purity 1.00 --tumor-purity 1.00 --p-value 0.99 --somatic-p-value 0.01 --strand-filter 0 --validation 1 --output-vcf 1”. Finally, fpfilter was performed with the configuration of “--min-var-freq 0.0025 --min-var-count 4 --min-strandedness 0.05 --min-ref-readpos 0.2 --min-var-readpos 0.2 --min-var-dist3 0.15 --min-ref-dist3 0.2 --min-var-basequal 30 --min-ref-basequal 30 --max-rl-diff 0.2 --max-mmqs-diff 1000000 --max-ref-mmqs 50 --min-ref-mapqual 20 --min-var-mapqual 30”.

### Performance evaluation metrics

Recall, precision and F1 score were calculated for performance evaluation:

recall = number of true positives / (number of true positives + number of false negatives), precision = number of true positives / (number of true positives + number of false positives), F1 score = 2*recall*precision / (recall + precision).

For model performance evaluation, receiver operating characteristic (ROC) curves were plotted and area under the curve (AUC) scores were calculated with R package PRROC.

### Development of SomaticXGB: Feature extraction, training, validation and optimization

Candidate variants were obtained from VarScan2 without fpfilter, which were input into force-calling mode of Mutect2 with the setting of “--genotype-filtered-alleles --alleles” to obtain some features for model training. Other features were obtained from BAM-readcount by inputting positions of candidate variants. Simulated variants were labeled as 1; other candidates were labeled as 0. After obtaining training data, R package XGBoost was used to train a model to score each candidate variant. Grid-search was performed to determine the best hyperparameter combination (max_depth of 3, 5, 7 and 9; eta of 0.01, 0.05, 0.1 and 0.2; nrounds of 100, 200 and 400). K-fold cross validation was conducted and AUC was used to evaluate the performance of each hyperparameter combination (Supplementary Fig. 3). The configuration of max_depth=7, nrounds=400 and eta=0.1 was the best and used to train the model.

### Calculations of error rates and VAFs

To calculate the error rates for HG001-HG005, we initially identified the regions by finding the overlap between our panel’s target regions and the high-confidence intervals provided by GIAB, and excluding the sites with germline mutations. Within these defined regions, the error rates were determined by calculating the proportion of mismatches to the total number of bases with a quality score of Q30 or higher. VAFs were calculated by dividing the allele depth by the total depth of coverage. Both the error rate and VAF calculations were carried out using an in-house program.

